# Fast surveillance response and genome sequencing reveal the circulation of a new Yellow Fever Virus sublineage in 2021, in Minas Gerais, Brazil

**DOI:** 10.1101/2021.11.24.469129

**Authors:** Miguel S. Andrade, Fabrício S. Campos, Cirilo H. de Oliveira, Ramon Silva Oliveira, Aline A. S. Campos, Marco A. B. Almeida, Danilo Simonini-Teixeira, Anaiá da P. Sevá, Andrea Oliveira Dias Temponi, Fernando Maria Magalhães, Danielle Costa Capistrano Chaves, Maira Alves Pereira, Ludmila Oliveira Lamounier, Givaldo Gomes de Menezes, Sandy Micaele Aquino Teixeira, Maria Eduarda Gonçalves dos Santos, Sofía Bernal-Valle, Nicolas F. D. Müller, Jader da C. Cardoso, Edmilson dos Santos, Maria A. Mares-Guia, George R. Albuquerque, Alessandro P. M. Romano, Ana C. Franco, Bergmann M. Ribeiro, Paulo M. Roehe, Filipe V. S. Abreu

**Affiliations:** Baculovirus Laboratory, Department of Cell Biology, Institute of Biological Sciences, University of Brasilia, Brasília, Distrito Federal, 70910-900, Brazil; Bioinformatics and Biotechnology Laboratory, Campus of Gurupi, Federal University of Tocantins, Gurupi, Tocantins, 77410-570, Brazil; Insect Behavior Laboratory, Federal Institute of Northern Minas Gerais, Salinas, Minas Gerais, 39560-000, Brazil; State Center of Health Surveillance, Rio Grande do Sul State Health Department, Porto Alegre, Rio Grande do Sul, 90610-000, Brazil; Department of Agricultural and Environmental Sciences, Santa Cruz State University, Ilhéus, Bahia, 45662-900, Brazil; State Arbovirus Surveillance Coordination, Minas Gerais State Health Department, Belo Horizonte, Minas Gerais, 31630-901, Brazil; Central Public Health Laboratory (LACEN-MG), Fundação Ezequiel Dias, Belo Horizonte, Minas Gerais, 30510-010, Brazil; Institute of Basic Health Sciences, Federal University of Rio Grande do Sul, Porto Alegre, Rio Grande do Sul, 90050-170, Brazil; Flavivirus Laboratory, Instituto Oswaldo Cruz, Fiocruz, Rio de Janeiro, Rio de Janeiro, 21040-360, Brazil; General Coordination of Arbovirus Surveillance, Ministry of Health, Brasília, Distrito Federal, 70058-900, Brazil

**Keywords:** Yellow fever virus, Arbovirus, Flavivirus, Non-human primate, Epizootic, Smartphone

## Abstract

Yellow fever virus (YFV) exhibits a sylvatic cycle of transmission involving wild mosquitoes and non-human primates (NHP). In Brazil, YFV is endemic in the Amazon region, from where waves of epidemic expansion towards other Brazilian states eventually occur. During such waves, the virus usually follows the route from North to the Central-West and Southeast Brazilian regions. Amidst these journeys, outbreaks of Yellow Fever (YF) in NHPs, with spillovers to humans have been observed. In the present work, we describe a surveillance effort encompassing the technology of smartphone applications and the coordinated action of several research institutions and health services that succeeded in the first confirmation of YFV in NHPs in the state of Minas Gerais (MG), Southeast region, in 2021, followed by genome sequencing in an interval of only ten days. Samples from two NHPs (one of the species *Alouatta caraya* in the municipality of Icaraí de Minas and the other of the species *Callithrix penicillata* in the municipality of Ubaí) were collected and the presence of YFV was confirmed by RT-qPCR. We generated three near-complete by Nanopore sequencer MinION. Phylogenetic analysis revealed that all viral genomes recovered are equal and related to lineage South America 1, clustering with a genome detected in the Amazon region (Pará state) in 2017. These findings reveal the occurrence of a new wave of viral expansion in MG, six years after the beginning of the major outbreak in the state, between 2015-2018. No human cases were reported to date, showing the importance of coordinated work between local surveillance based on available technologies and support laboratories to ensure a quick response and implementation of contingency measures towards avoiding the occurrence of YF cases in humans.

## 1) Introduction

In Brazil, the yellow fever virus (YFV, family *Flaviviridae*, genus *Flavivirus*) exhibited two epidemiologically distinct transmission cycles, urban and sylvatic, although the former must be mentioned here from a historical perspective. In the urban cycle, which has not been recorded since 1942, the virus is transmitted among humans by the vector *Aedes aegypti* (Franco, 1969; Monath and Vasconcelos, 2015). In the sylvatic/jungle cycle, the virus is transmitted by wild mosquitoes (mainly *Haemagogus* and *Sabethes*) to non-human primates (NHPs, e.g. *Alouatta* and *Callithrix* genera) and, occasionally, to unvaccinated humans in close contact with forest areas. Yellow fever (YF) is considered endemic in the tropical rainforest of the Amazon region, from where waves of epidemic expansion spread towards other Brazilian regions at irregular intervals of time (Possas et al., 2018). During these waves, the virus usually reaches the states of Goiás (Central-West) and Minas Gerais (Southeast region). Sometimes, the virus arrives in other southeast states, such as São Paulo (in the years 2000, 2008-2009, 2017-2018) (Vasconcelos et al., 2001; Camargo-Neves et al., 2005; de Souza et al,. 2011; Moreno et al., 2011; Romano et al., 2014; Moreno et al., 2015; Cunha et al., 2019), Rio de Janeiro and Espírito Santo (between the years 2017 and 2019) (Fernandes et al., 2017; Gómez et al., 2018; Abreu et al., 2019b; Delatorre et al., 2019), and even the southernmost states of the country, Rio Grande do Sul (2001, 2008-2009, 2020-2021) (Almeida et al., 2012; Vasconcelos et al., 2003; Andrade et al., 2021), Paraná e Santa Catarina (2018-2020) (Ministry of Health Brazil, 2021). The expansion of YF outside the endemic areas raises, at least, three main concerns: 1) the possibility of re-emergence of an urban cycle; 2) the increased risk of YF in humans due to heterogeneous vaccination coverage outside the Amazon region; 3) the risk of extinction of threatened NHP species, especially from the genus *Alouatta*, highly susceptible to YFV (Holzmann et al., 2010; Almeida et al., 2012; Romano et al., 2014; Bicca-Marques et al., 2017; Dietz et al., 2019; Strier et al., 2019; Berthet et al., 2021).

During the expansion waves, the state of Minas Gerais (MG) has been an important corridor through which the virus travels before spreading to other Brazilian states (Delatorre et al. 2019). In recent times, major outbreaks occurred in MG in 2000-2003, 2010, and between 2015-2018 (Fig. 1A), the latter being the largest sylvatic outbreak in the last 80 years (Causey et al., 1949; Vasconcelos et al., 2001; Costa, 2005; Ribeiro and Antunes, 2009; Pinheiro et al., 2019; Cunha et al., 2019). In the 2015-2018 outbreak, two viral sub-lineages were detected, crossed MG by different paths: the YFV_MG/SP/RS_ sub-lineage traveled through the West and Southwest of the state, before reaching the state of São Paulo; the YFV_MG/ES/RJ/BA_ sub-lineage, spread by the Northwest, North, Northeast, and East of the state, before reaching the states of Espírito Santo and Bahia (Delatorre et al., 2019; Goes de Jesus et al., 2020; Andrade et al., 2021). Although epizootics have been constantly reported since the end of 2020 (without the opportunity for sample collection), the last reported viral detection in MG occurred in the first semester of 2018 (Delatorre et al., 2019; Minas Gerais, 2021).

**Figure 1.**
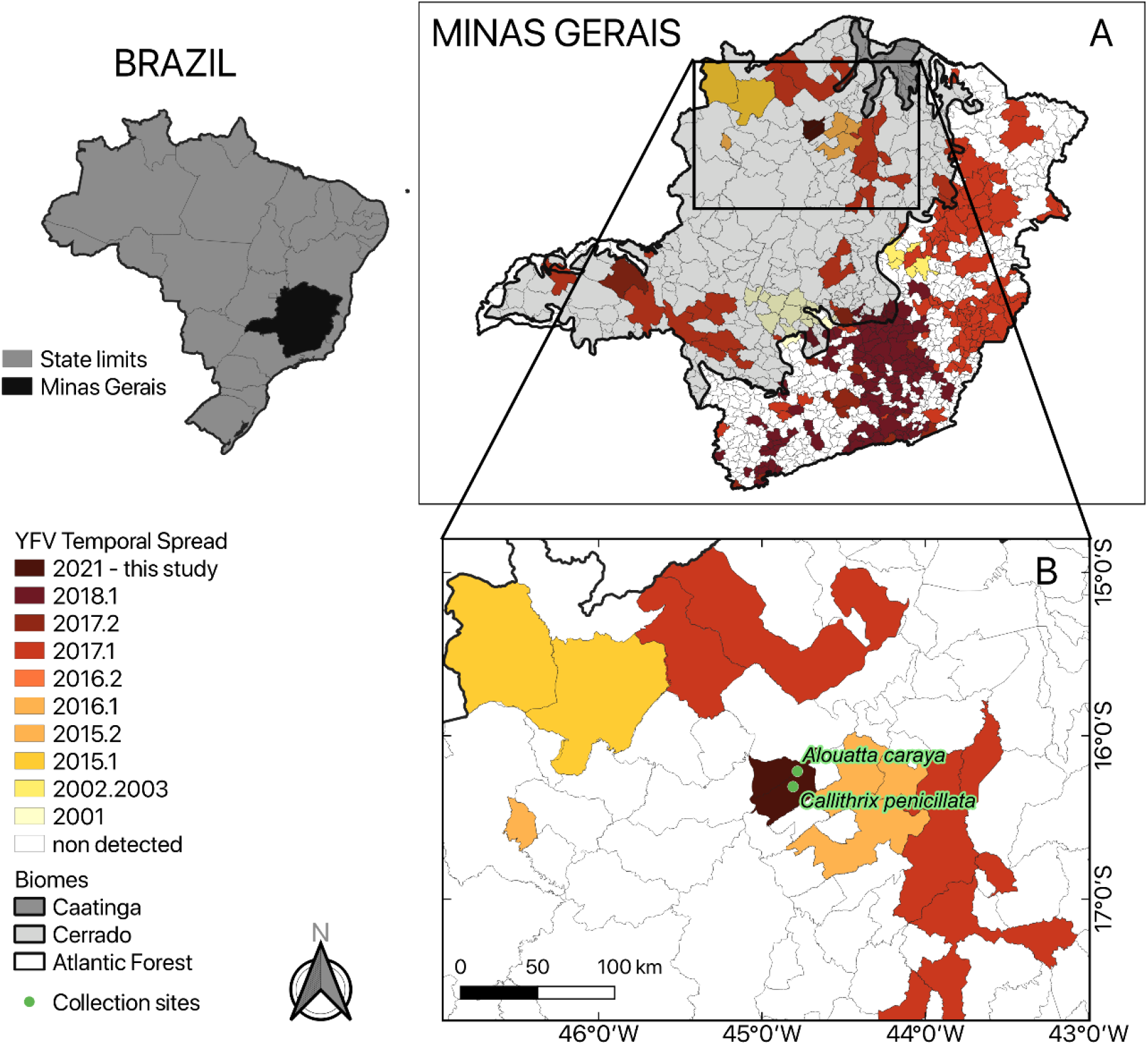
On the left, the Brazil map highlights Minas Gerais (MG) state. The Brazilian regional division consists of states and municipalities grouped into regions. A) MG state map showing different municipalities with YFV detections in NHP and/or humans between 2001 to present (see the caption on the left). In gray are shown the different biomes of MG (Caatinga, Cerrado, and Atlantic Forest). B) Location of epizootics in Minas Gerais state registered on the SISS-Geo platform during this investigation.

In this work, we describe a surveillance effort encompassing the technology of smartphone applications and the coordinated action of State Arbovirus Surveillance Coordination of Minas Gerais State Health Department, Central Public Health Laboratory (LACEN-MG), Ministry of Health of Brazil, and members of Project “Febre Amarela BR” (Portuguese for “Yellow Fever BR”) that succeeded in the first confirmation of YFV in NHPs in the state of Minas Gerais (MG) in 2021, followed by genome sequencing, in an interval of only ten days. Anticipating our findings, phylogenetics analysis reveals the introduction of a new sub-lineage in the extra Amazonian region.

## 2) Material and Methods

### 2.1) Study area

This study was carried out in the state of Minas Gerais, which concentrates the largest number of municipalities (853) in Brazil, the second largest population in the country (21,411,923 inhabitants or 10.1% of the Brazilian population), and the fourth largest geographic extension (586,513.993 km2, larger than Spain, for example) (https://www.ibge.gov.br/cidades-e-estados/mg.html). The area covered by this study belongs to the North region of Minas Gerais State, which is predominantly covered by Cerrado (a savannah-like biome) (**Figure 1**). The region has well-defined dry and rainy periods, with December presenting the highest average rainfall during the summer, and August, the month of this investigation, the driest month of the year, during the winter.

### 2.2) Building an information network to strengthen epizootic surveillance in northern Minas Gerais

Since 2020, we have started the organization of an information network linked to a research project called Febre Amarela BR (Yellow fever BR), with the objective of aggregating research institutions, municipal, state, and federal health agencies, in addition to the general population, to strengthen the surveillance of epizootics (Abreu et al., 2019a; Abreu et al., 2019c).

For these purposes, we organized preliminary field expeditions to several municipalities in Minas Gerais. In each municipality, meetings and lectures were organized with health agents and environmental surveillance agents, aiming to inform the teams about the importance of epizootics surveillance. The agents were trained to notify epizootics using the SISS-Geo web app (Chame et al., 2019). SISS-Geo is the abbreviation of “Sistema de Informação em Saúde Silvestre Georreferenciado” in Portuguese or “Georeferenced Wildlife Health Information System”, which is a platform that works as a public repository for collaborative surveillance of wild animals in Brazil, for the registration of animals occurrences sending pictures of dead or alive animals, and also metadata. The app collects real-time geographic coordinates and sends the information to state and federal surveillance centers. We also organized field works to train the teams to perform vectors and NHP sample collection. Finally, several groups were created in the WhatsApp application with all municipal health agents reached through the meetings, constituting an information network, to send educational material about vector-borne diseases and to exchange news about epizootics (Abreu et al., 2019c). An organization chart showing the steps for organizing the network is shown in **Figure 2**. All field expeditions were carried out within the scope of the Yellow Fever BR project, with the support of the State Health Department and its regional coordinators.

**Figure 2.**
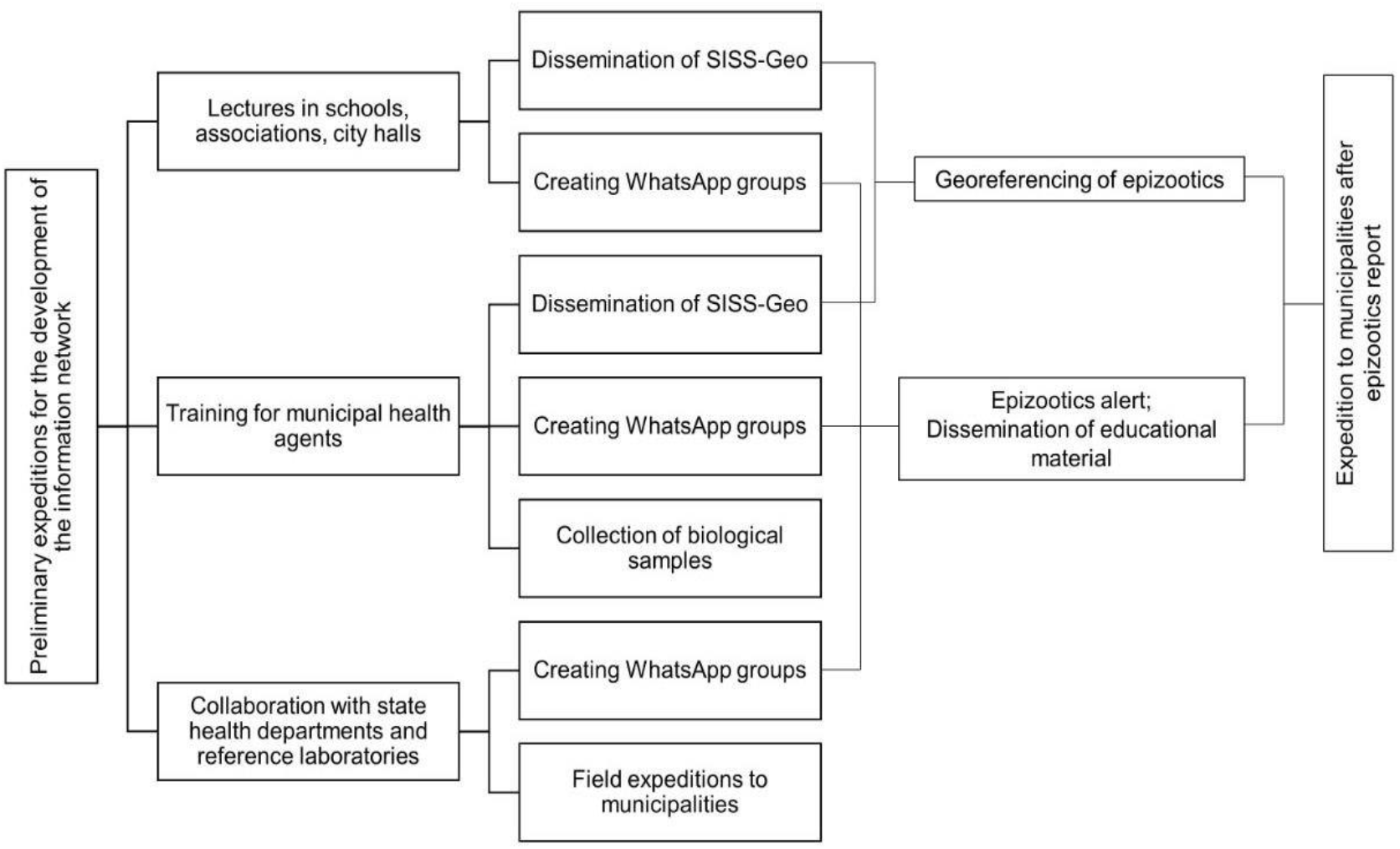
Organization chart showing steps for organizing the information network to strengthen yellow fever surveillance

### 2.3) Investigation of epizootic in Northern MG

On August 24, 2021, the information network alerted us about the occurrence of epizootics in the municipalities of Icaraí de Minas and Ubaí, located in the northern region of Minas Gerais, 595 km from Belo Horizonte, the capital of the state. Interestingly, these municipalities were not affected by the 2015-2018 outbreak. Focusing on a timely sampling, an expedition was rapidly organized on the following morning, August 25th, 2021, composed of members of the Febre Amarela BR project, members of the state health department, and agents from the affected municipalities. On August 25th and 26th of 2021, we surveyed the riparian forests of the places searching NHP carcasses for the collection of biological samples. The detected epizootics were properly registered in the SISS-Geo app and in the Brazilian Information System on Notifiable Diseases (SINAN). Samples were collected following safety protocols (Brasil, 2017), preserved in liquid nitrogen (−196°C) and sent to the reference laboratory of the national public health system, whose destination in Minas Gerais was the Ezequiel Dias Foundation (FUNED) as well as to the Baculovirus Laboratory, a branch of the research project Febre Amarela BR for viral diagnosis, at University of Brasilia.

### 2.4) Molecular diagnosis

Samples (different viscera fragments) were lysed with Trizol™, stored in liquid nitrogen and sent to the sequencing laboratory in dry ice (−80°C). Total RNA extraction with Trizol™ was performed following the manufacturer’s instructions. YFV RNA detection was done using two previously published RT-qPCR protocols (Domingo et al., 2012).

### 2.5) Genome sequencing

Samples with CT <25 were submitted to cDNA synthesis protocol using LunaScript™ RT SuperMix Kit (NEB) following the manufacturer’s instructions. Then, a multiplex tiling PCR was performed using the previously published YFV primers (Faria et al., 2018) and 40 cycles (denaturation: 95°C/15 s and annealing/extension: 65°C/5 min) of PCR using Q5 high-fidelity DNA polymerase (NEB). Amplicons were purified using 1× AMPure XP beads (Beckman Coulter), and cleaned-up PCR product concentrations were measured using a QuantiFluor^®^ dsDNA System assay kit on a Quantus™ Fluorometer (Promega). DNA library preparation was performed using the Ligation sequencing kit SQK-LSK309 (Oxford Nanopore Technologies) and the Native barcoding kit (EXP-NBD104 and EXP-NBD114; Oxford Nanopore Technologies, Oxford, UK). The sequencing library (23 samples and a negative control per run) was loaded onto an R9.4 flow cell (Oxford Nanopore Technologies) and sequenced between 6 to 18 hours using MiNKOW software. The RAMPART (Version 1.2.0, ARTIC Network) package was used to monitor coverage depth and genome completion. The resulting Fast5 files were basecalled and demultiplexed using Guppy (Version 4.4.2, Oxford Nanopore Technologies). Variant calling and consensus genome assembly were carried out with Medaka (Version 1.0.3, Oxford Nanopore Technologies) using the sequence JF912190 as the reference genome.

### 2.6) Phylogenetic analyses

To perform phylogenetic analyses, we selected all near-complete YFV sequences from NCBI (n=381, excluding sequences < 8 kb and those of vaccine and patent-related viruses). Metadata as samples collection date and geographic coordinates were retrieved from GenBank files or from genome associated publications (manual curation). Genomes MG67-L combined with the 381 genomes from NCBI were aligned with MAFFT v.7.480 (Katoh and Standley, 2013). The Maximum-likelihood tree was inferred using IQTREE, with the GTR+F+I+┌4 model. The new genome sequences were sent to the NCBI GenBank database under accession numbers OL519587 to OL519589.

## 3) Results

On August 25, 2021 (day 1), we notified an epizootic affecting six black-and-gold howler monkeys (*Alouatta caraya*) in Icaraí de Minas, of which only one was suitable for sampling (liver, spleen, kidney samples were collected and named MG66-L, MG66-S, and MG66-K, respectively). On August 26 (day 2), another epizootic was detected, affecting six black-pincelled marmosets (*Callithrix penicillata*) in Ubaí and we were able to collect samples from one (liver, spleen, kidney and brain, named MG67-L, MG67-S, MG67-k and MG67-B, respectively). Samples were sent to the laboratory (September 02, day 9) and tested (September 03, day 10). All tissues sampled were positive for YFV by RT-qPCR (**Table 1**).

**Table 1.** RT-qPCR test results, according to tested species, local, tissues, date, and Ct.

| Host species | Municipality | Geographical coordinates | Sample | Collection date | Ct |
| --- | --- | --- | --- | --- | --- |
| *Alouatta caraya*<br>(MG66) | Icaraí de Minas | 16°13'03.3"S<br>44°47'00.9"W | Liver | 25/08/2021 | 15 |
|  |  |  | Spleen | 25/08/2021 | 14 |
|  |  |  | Kidney | 25/08/2021 | 20 |
| *Callithrix penicillata*<br>(MG67) | Ubaí | 16°18'42.0"S<br>44°48'36.0"W | Liver | 26/08/2021 | 31 |
|  |  |  | Spleen | 26/08/2021 | 30 |
|  |  |  | Kidney | 26/08/2021 | 31 |
|  |  |  | Brain | 26/08/2021 | 29 |

The interval between the alert of our information network and diagnosis was 10 days (**Figure 3**). After confirming the detection of the virus, health authorities were immediately communicated. Intensified control and investigation actions in the affected and surrounding areas were implemented. Meanwhile, the sequencing library was prepared on the same day and applied to the sequencer resulting in an 8-hour interval between RNA extraction and first read generated.

**Figure 3.**
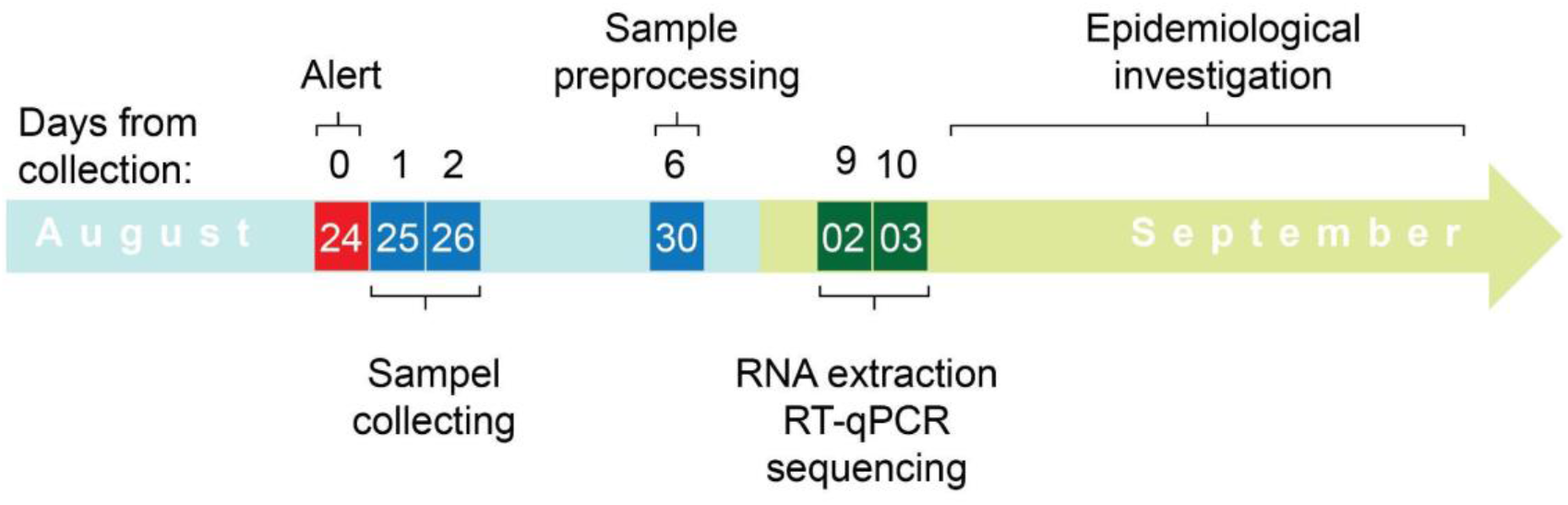
Timeline showing the time spent from the collection of samples until the generation of near-complete genomes.

Three near-complete YFV genomes were generated from different viscera of the same animal: MG66-Liver, MG66-Spleen, and MG66-kidney. The three YFV genomes of *Alouatta caraya* (MG66) were compared and revealed no intra-host sequence differences. Therefore, for phylogenetic analyses, the MG66-L genome was used.

Phylogenetic analysis revealed that the YFV genomes generated here were related to the lineage South America 1, as expected, and clustered with an isolate from NHP (*Alouatta caraya*) identified in Pará (Amazonian region) in 2017 (**Figure 4**) showing that they are not related to the viruses detected on this same year (2021) in the state of Rio Grande do Sul, the southernmost state of Brazil (Andrade, 2021). These results indicate the circulation of at least two sub-lineages of YFV in the extra amazon region in 2021.

**Figure 4.**
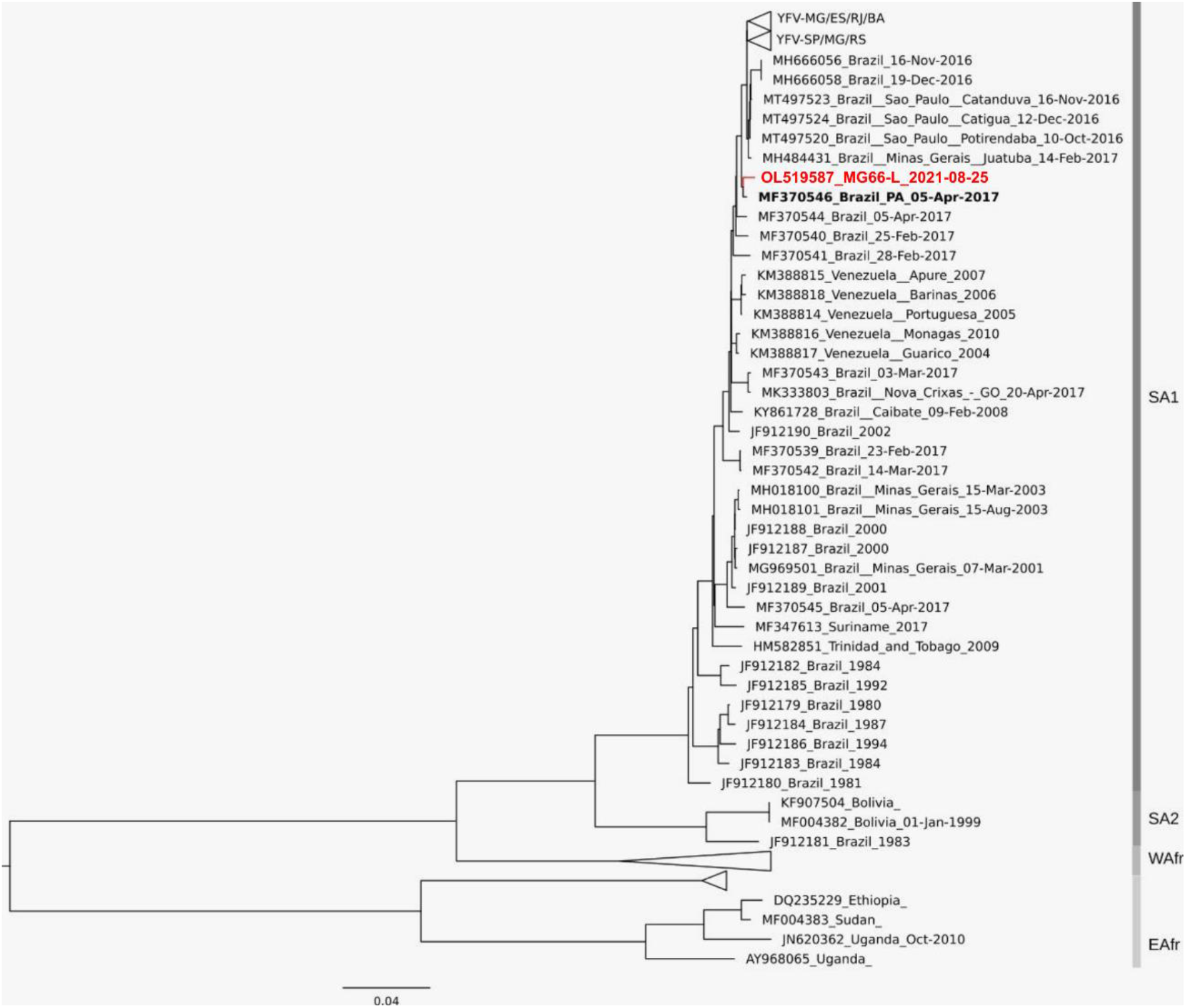
Phylogenetic tree of YFV based on MG66-Liver and NCBI genomes. The collapsed group includes YFV_MG/ES/RJ/BA_ and YFV_MG/SP/RS_. South America I (SAI), South America 2 (SA2), West Africa (WAfr), and East Africa (EAfr) genotypes are indicated. YFV MG/2021 is highlighted in red and is related to the lineage SA1, the same present in the Amazon region in 2017. The sub-lineages that caused the 2015-2021 outbreak (YFV_MG/SP/RS_ and YFV_MG/ES/RJ/BA_) are shown at the top of the tree.

## 3) Discussion

The state of Minas Gerais has been historically affected by several introductions of the YFV. However, logistical limitations make it difficult to carry out surveillance and the timely collection of biological samples, which requires a robust field and laboratory surveillance system. In the present work, we describe a result of the construction of an inter-institutional network capable of collaborative actions to investigate epizootics, from detection and sample collection to genomic sequencing. The rapid outbreak response (only 10 days between the alert of an epizootic and genome sequencing) carried out in August/2021 resulted in the first confirmation of the circulation of YFV in the state of Minas Gerais in 2021 and showed that the recovered viral genome belongs to the sub-lineage that circulated in Pará in 2017 (now called YFV_PA/MG_).

The state of Minas Gerais has played a role as a YFV spreading route during the expansion waves from the Amazon region. Interestingly, there have been studies of viral spread routes in the region since 1938-1943 (**Suppl. Fig. 1**) (Causey et al., 1949).

In the last two decades, outbreaks were recorded in 2001 (32 human cases), 2002 to 2003, (63 human cases), and two isolated cases between 2008-2009. However, the largest outbreak was recorded between 2016-2018, with 1006 human cases, and 448 confirmed epizootics (Costa, 2005; Ribeiro and Antunes, 2009; Minas Gerais, 2018; Minas Gerais, 2019). Importantly, there were no records of previous viral circulation in the two currently affected municipalities, which may indicate the presence of a naive NHP population (Fernandes et al., 2021; Mares-Guia et al., 2020).

Since 2019, as far as we know, despite the notification of several epizootics, no new confirmations of YFV circulation occurred in the north of MG (Minas Gerais, 2021). The wide territorial extension, with huge rural areas far from urban and research centers, combined with socioeconomic inequality and the lack of trained professionals in several municipalities make it difficult to implement agile and uniform surveillance of epizootics, diminishing the chances of timely sample collection for virus detection. These characteristics reinforce the need to use new tools and strategies to strengthen surveillance. Among them, worth it to mention: a) the use of smartphone applications to notify epizootics and collect the geographic coordinates of the occurrence in real time. In this sense, the SISS-Geo app, which has been progressively implemented in Brazil, speeds up the arrival of information to involved institutions and provides accurate geographic location, streamlining outbreak response (Chame et al., 2019); b) the use of messaging apps such as Whatsapp to increase the number of “watchers”. Creating Whatsapp groups with local or regional reach, composed by different members of the population (health agents, cyclists, hikers, students, rural workers, etc.) increases the chances of detecting an epizootic and speeds up the dissemination of important information such as vaccination campaigns and monkey conservation efforts (Abreu et al., 2019 b and c); c) creation of multi-institutional networks to facilitate and speed up sample collection and diagnosis. In the present work, research institutions linked to the Febre Amarela BR project collaborated with municipal, state, and federal health secretariats. With this joint effort, in the space of 10 days, it was possible to detect and notify the epizootic, collect samples in a distant and isolated region, carry out the molecular diagnosis, disseminate the results to the local governments, and generate the near complete genome of the virus and its phylogeographic relationship (called real-time genomic surveillance). Thus, measures to protect the population, such as the expansion of vaccination coverage, urban vector control, and educational campaigns were triggered immediately by Minas Gerais health authorities, mitigating the risk of YF human cases.

The implementation of real-time genomic surveillance showed that the virus circulating in Minas Gerais is related to the lineage that circulated in the state of Pará, in the Amazon region, in 2017. Therefore, it is a new wave of viral expansion in MG, six years after detection in 2015, when the virus actively spread through the state until 2018. Interestingly, in 2021, at least two different viral sub-lineages circulating at the same time in the extra-Amazon region, the YFV_MG/SP/RS_, detected in the Rio Grande do Sul (Andrade et al., 2021), and YFV_PA/MG_, described in the present work, in Minas Gerais. And it is noteworthy that the virus was detected at the height of the dry season, and in one of the driest regions of MG, which makes it difficult for the main species of mosquito vectors to survive.

Here, in this brief report, we can see the importance of coordinated work between the municipal health secretariats, the state health secretariat, and support laboratories. Through this system, it was possible to collect the PNH samples, process them and quickly give a result for YFV enabling the taking of protective measures by the competent agents. The genomic surveillance showed that it is a new wave of expansion of the YFV in Minas Gerais, which must be monitored in order to improve preventive actions such as elevating the vaccine coverage in humans and improve communication about the YFV risk to health professionals and the general population, in order to avoid human cases in other areas of Minas Gerais and Brazil. Also, this should improve the sensibilization about the importance of the non-human-primates as sentinels as for YFV presence in the region. Finally, this study shows the importance of collaborative, integrated surveillance, allowing prompt action, even in challenging situations, with stakeholders able to support and answer time on the yellow fever surveillance and response.

## Author Contributions

Conceived and designed the experiments: F.V.S.A, M.S.A., F.S.C., A.A.S.C., J.C.C., E.S., D.S.T., G.R.A., A.C.F., B.M.R., P.M.R. and M.A.B.A. Performed the experiments: M.S.A., M.A.P., L.O.M, S.M.A.T, M.E.G.S, F.L.M., J.C.C., E.S., L.C.B., C.M.D.S., N.F.D.M., C.H.O., A.J.J.S., S.B.V., M.A.M.G., G.G.M., L.O.M and M.A.B.A. Analyzed the data: M.S.A., F.S.C., A.A.S.C., F.V.S.A., F.L.M., A.P.S., J.C.C., E.S., D.S.T., A.P.M.R., B.M.R., A.O.D.T, P.M.R., and M.A.B.A. Contributed reagents/materials/analysis tools: A.C.F., B.M.R., P.M.R., A.O.D.T, D.C.C.C and M.A.B.A. Contributed to the writing of the manuscript: M.S.A., F.S.C., A.A.S.C., M.A.B.A., F.L.M., A.P.S., J.C.C., E.S., A.O.D.T., D.S.T., S.B.V., A.P.M.R., B.M.R., P.M.R. and F.V.S.A.

## Acknowledgments

We acknowledge the contributions of the Division of Environmental Health Surveillance from Minas Gerais State. The authors would like to thank the effort of the MG Yellow Fever Surveillance Team that was at the forefront of the preparation to face the arrival of the virus in the state as well as in field investigations. To Beatrízio Rodrigues Almeida, Hermes Almeida and Joselio Ribeiro Paraíso for their valuable help during the field work. To Aline Tátila Ferreira for her help in creating images. We are also grateful to countless colleagues from municipalities’ health departments, who conducted the investigation of epizootics collecting samples in the field and to the Ministry of Health’s Arbovirus Surveillance Team. The Yellow Fever Brazil project (Febre Amarela BR: https://www.febreamarelabr.com.br/) is supported by grants from Conselho Nacional de Desenvolvimento Científico e Tecnológico and Departamento de Ciência e Tecnologia of Secretaria de Ciência, Tecnologia e Insumos Estratégicos of Ministério da Saúde (CNPq/Decit/SCTIE/MS grant number 443215/2019-7). M.S.A. is granted a post-doctoral scholarship (DTI-A) from CNPq. P.M.R. is a CNPq research fellow.

