## Supplementary material for "Fast surveillance response and genome sequencing reveal the circulation of a new Yellow Fever Virus sublineage in 2021, in Minas Gerais, Brazil": Suppl. Fig. 1

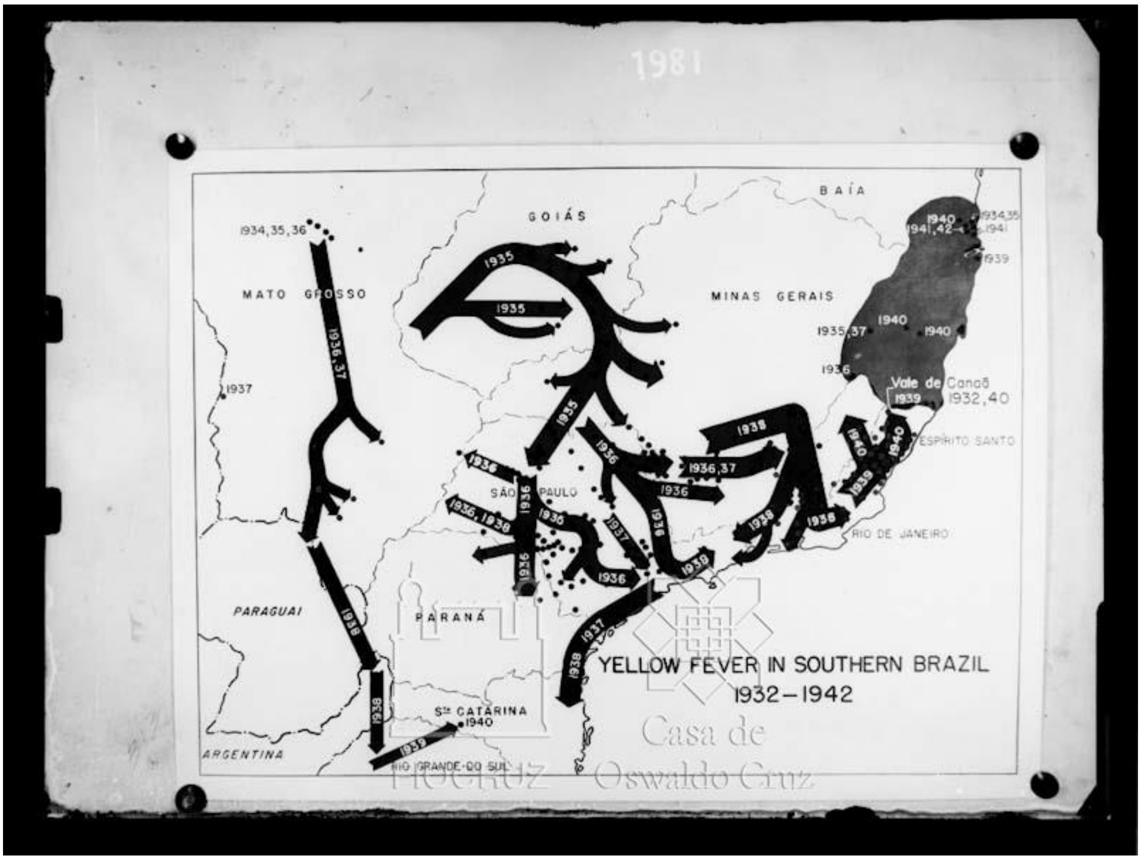


**Supplementary Figure 1**. Map demonstrating the spread of yellow fever in southern Brazil, 1932-1942 Source: Casa de Oswaldo Cruz Collection, Department of Archives and Documentation - Photo FR (SFA-EC) 12-5 by A. Fialho (http://arch.coc.fiocruz.br/index.php/y0luj).
